# DeepInsight-3D for precision oncology: an improved anti-cancer drug response prediction from high-dimensional multi-omics data with convolutional neural networks

**DOI:** 10.1101/2022.07.14.500140

**Authors:** Alok Sharma, Artem Lysenko, Keith A Boroevich, Tatsuhiko Tsunoda

**Affiliations:** Laboratory for Medical Science Mathematics, RIKEN Center for Integrative Medical Sciences, Japan; Institute for Integrated and Intelligent Systems, Griffith University, Australia; Laboratory for Medical Science Mathematics, Department of Biological Sciences, School of Science, The University of Tokyo, Tokyo, Japan; Laboratory for Medical Science Mathematics, Department of Computational Biology and Medical Sciences, Graduate School of Frontier Sciences, The University of Tokyo, Japan

## Abstract

Modern oncology offers a wide range of treatments and therefore choosing the best option for particular patient is very important for optimal outcomes. Multi-omics profiling in combination with AI-based predictive models have great potential for streamlining these treatment decisions. However, these encouraging developments continue to be hampered by very high dimensionality of the datasets in combination with insufficiently large numbers of annotated samples. In this study, we propose a novel deep learning-based method to predict patient-specific anticancer drug response from three types of multiomics data. The proposed DeepInsight-3D approach relies on structured data-to-image conversion that then allows use of convolutional neural networks, which are particularly robust to high dimensionality of the inputs while retaining capabilities to model highly complex relationships between variables. Of particular note, we demonstrate that in this formalism additional channels of an image can be effectively used to accommodate data from different ‘omics layers while explicitly encoding the connection between them. DeepInsight-3D was able to outperform two other state-of-the-art methods proposed for this task. These advances can facilitate the development of better personalized treatment strategies for different cancers in the future.

## Introduction

Precision oncology is rapidly developing. However, only a very small percentage of patients can currently take advantage of it [1]. The risks of side effects can be reduced by improving the prediction rate of drug response from targeted therapy which would undoubtedly improve patients’ treatment. In this respect, *in vitro* projects have compiled datasets such as Genomics of Drug Sensitivity in Cancer (GDSC) [2] and Cancer Cell Line Encyclopedia (CCLE) [3]. These datasets consist of multi-omics profiles such as gene expression, copy number alteration (CNA) and somatic mutations. Although gene expression datasets have shown to be very useful [2, 4], adding more omic layers could improve the predictability of pan-cancer models [5].

Drug prediction on *in vivo* datasets is a step towards clinical applicability. However, since *in vivo* data like The Cancer Genome Atlas (TCGA) repository has a scarcity of patient records and drug responses, it is difficult to train a model on *in vivo* datasets. Considering this challenge, Sharifi-Noghabi et al. [5] trained their computational model on *in vitro* data and obtained predictability on *in vivo* data. Nevertheless, the platforms of these datasets are different. Furthermore, there exists a scarcity of related samples in GDSC datasets. This augments the problem of adequately estimating a model on the training set of the data.

In [5], the MOLI method is proposed and learned on datasets from GDSC. It predicts response to a drug from TCGA and patient-derived xenograft (PDX) [6] datasets. These collated datasets have multi-omics profiles: gene expression, CNA, and somatic mutation. The MOLI method reported the area under the curve (AUC) on seven datasets from TCGA and PDX. Their average AUC over the seven datasets was 0.63. Park et al. [7] also constructed training and test dataset from GDSC and TCGA/PDX resulting in more than seven datasets and proposed the Super.FELT method. The performance of their method also included the seven datasets used in [5], though some of the resulting test samples slightly differed due to methodology. Nonetheless, their average AUC over the seven datasets was reported to be 0.68. When we reimplemented the Super.FELT method on the MOLI test sets, an average AUC of 0.65 was obtained.

In this work, we propose the DeepInsight-3D model, which first converts multi-omic or multi-layered data into corresponding images and then applies a convolutional neural network (CNN) to find the AUC. While the sample sizes of datasets are minimal, DeepInsight-3D with CNN can still perform well if the model is appropriately tuned. To compare with the benchmarked methods, we also utilized the same seven test sets as the MOLI study and achieved an average AUC of 0.72.

The proposed method is based on the DeepInsight method [8] and the DeepFeature method [9]. The DeepInsight method pioneered a strategy by converting non-image data to image form and then processing it to CNN for classification for various kinds of datasets. It has been widely used in various fields such as in cancer research [10-12], viral infections [13], sparse data [14], power energy [15], business and manufacturing [16], time-series data [17-19], traffic cash analysis [20], human activity recognition [21], feature representation [22], intrusion detection [23], spine surgery [24] and HVAC fault diagnosis [25]. Moreover, DeepInsight was a component in the Kaggle.com competition hosted by MIT and Harvard University that secured rank1 on the leaderboard [26]. Using dimensionality reduction techniques, such as t-SNE [27], DeepInsight arranges similar elements together in a 2D pixel frame and then performs element mappings. It turns tabular data into organized images, allowing CNN classification through automatic feature extraction. Furthermore, the DeepFeature method applied class-activation maps (CAMs) [28] to perform feature selection.

The DeepInsight-3D model extends the utility of DeepInsight and DeepFeature methods to multi-omics data, particularly 3D layers. It is well known that if the samples are sufficiently large, then CNN performs very well. Nonetheless, we have shown that the DeepInsight-3D model can also perform on a limited sample case. The overview of the proposed model is given in Fig. 1 (see Methods for the details).

**Figure 1.**
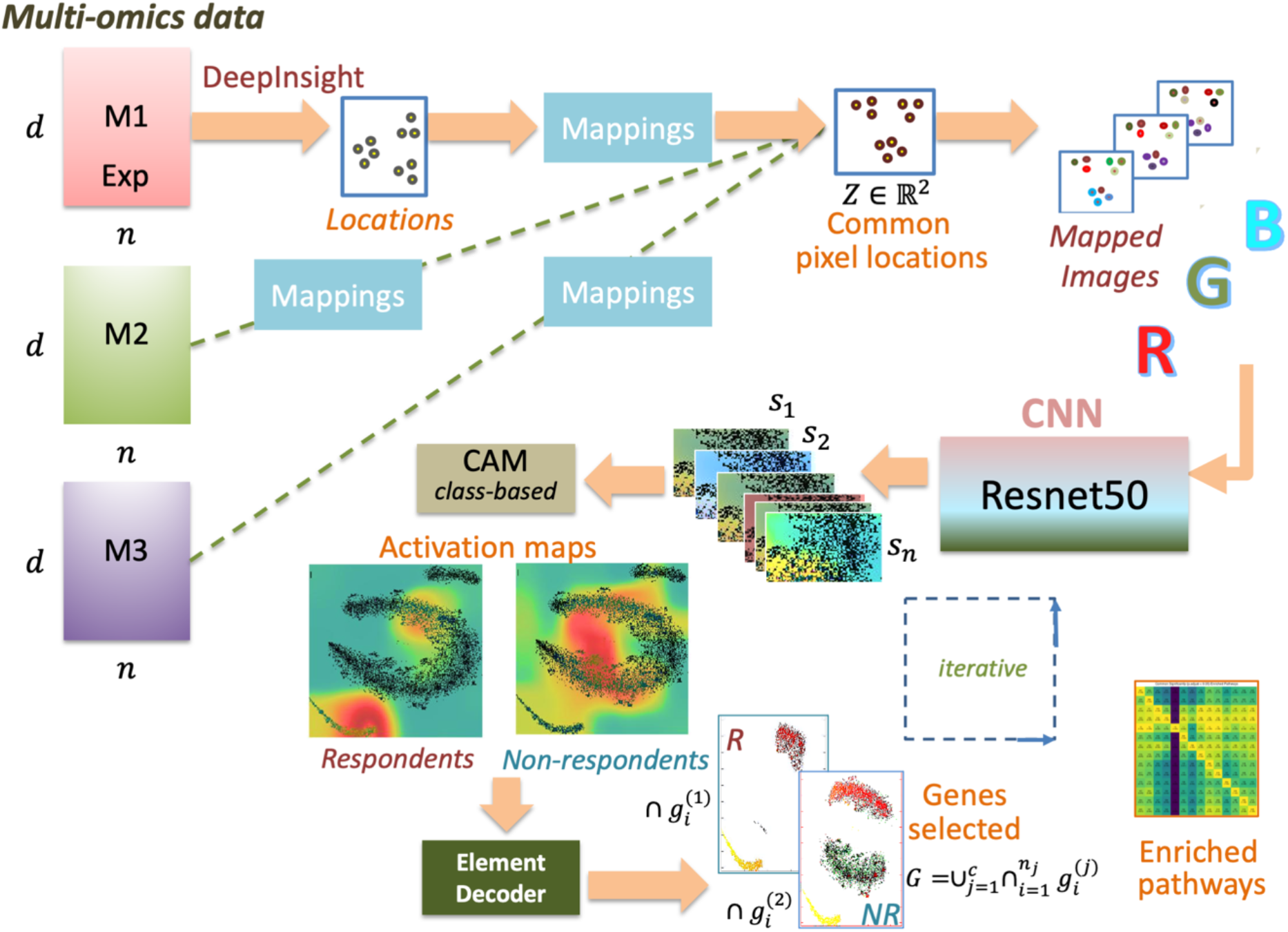
An overview of the DeepInsight-3D model. From the left multi-omics layers are processed via DeepInsight methodology and common pixel locations are found. After mapping omics data, corresponding images are constructed, which are processed to a convolutional neural network. Afterwards, CAM is used to find activation regions and element decoder is used to find a subset of genes.

In this work, DeepInsight-3D is used for multi-omics datasets. However, the proposed method is not limited to omics data. It can handle different kinds of multi-layered tabular data (as long as the elements and samples of diverse layers are arranged in the same order). This method does not require any specific biological information such as chromosome locations and visualizes non-image data through multi-layered mappings.

The contributions of this work are as follows. DeepInsight-3D pipeline is presented where classification and feature selection can be performed for multi-layered non-image samples (or tabular data) through the application of CNNs. Two ways of image construction are introduced, 1) by mapping elements to the pixel locations of the dominant layer (shown in Fig. 1), and 2) by mapping elements to the pixel locations obtained by giving equal importance to all the three layers (implemented in the DeepInsight-3D package as an option). Element decoder is implemented to find genes or elements from the activation maps. We also demonstrate how the developed system can be used to interpret the CNN model and report on the identified key genes and biological processes identified as important for drug response prediction by respective models.

## Results

### Performance evaluation

Two recently developed methods, MOLI and Super.FELT, were used as benchmark methods. Both these methods were compared with many preceding algorithms and showed superior performance. The test set configurations (in terms of the number of samples) were kept the same for a fairer comparison. The AUCs were computed for all possible drugs-method combinations and are given in Table 2.

**Table 1:**
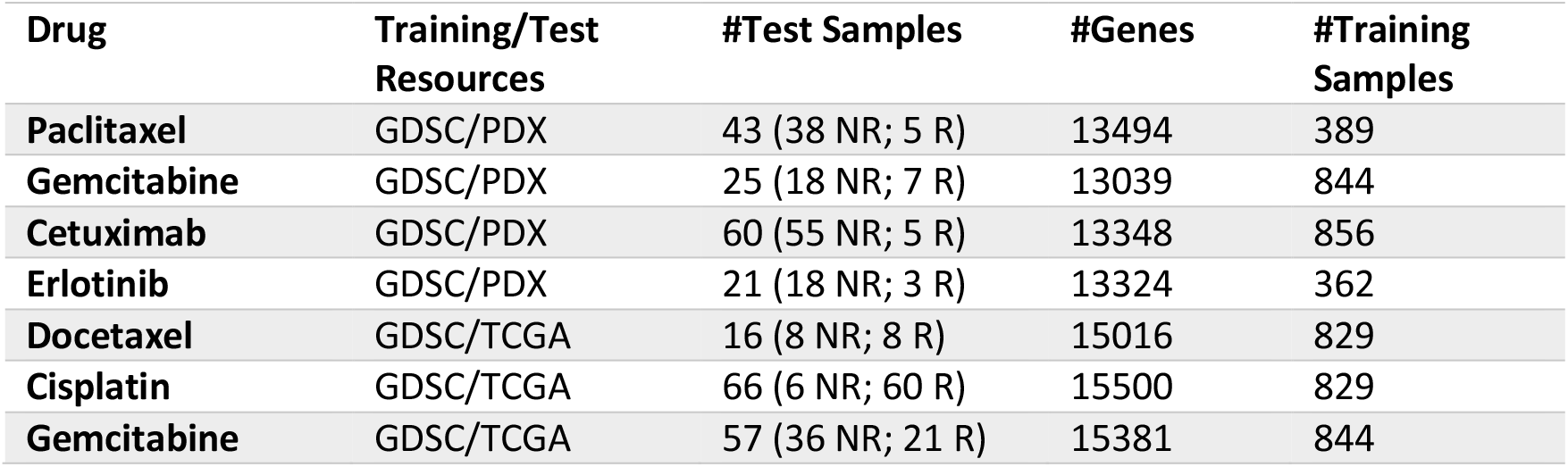
Training and test sets configurations for the drugs with multi-omics profiles.

**Table 2:**
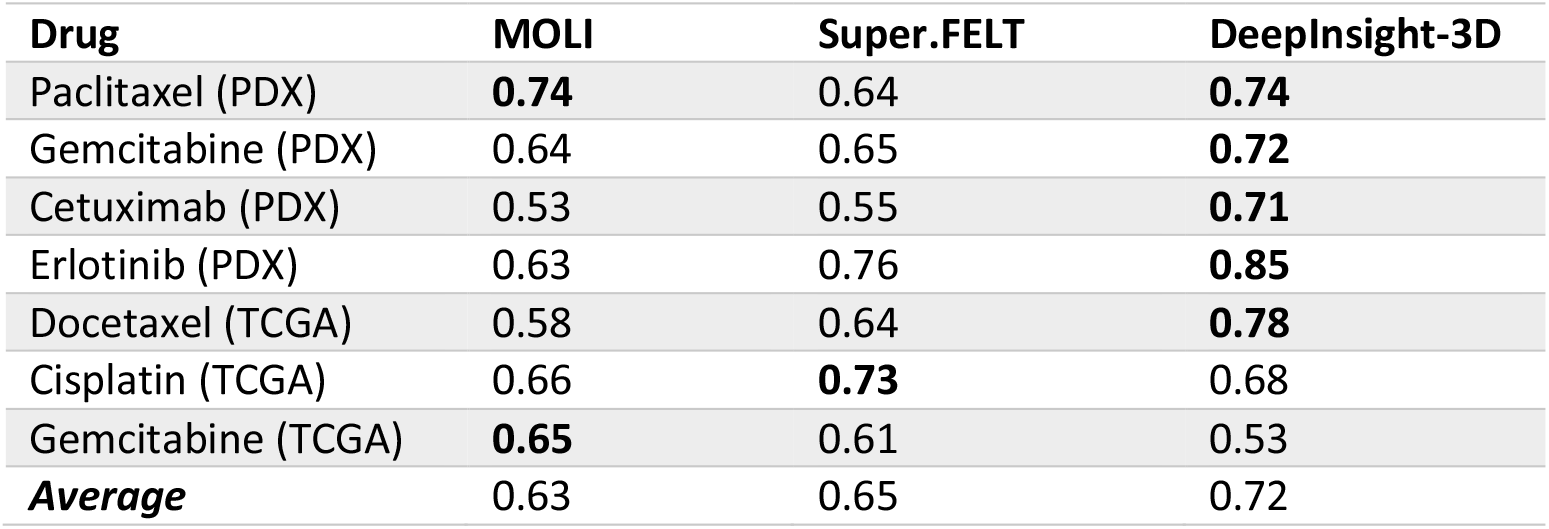
A comparison of DeepInsight-3D for drug response prediction with multi-omics profiles using test AUCs. The test samples are the same for all the comparators. The highest results are marked with bold fonts.

| Drug | MOLI | Super.FELT | DeepInsight-3D |
| --- | --- | --- | --- |
| Paclitaxel (PDX) | **0.74** | 0.64 | **0.74** |
| Gemcitabine (PDX) | 0.64 | 0.65 | **0.72** |
| Cetuximab (PDX) | 0.53 | 0.55 | **0.71** |
| Erlotinib (PDX) | 0.63 | 0.76 | **0.85** |
| Docetaxel (TCGA) | 0.58 | 0.64 | **0.78** |
| Cisplatin (TCGA) | 0.66 | **0.73** | 0.68 |
| Gemcitabine (TCGA) | **0.65** | 0.61 | 0.53 |
| **Average** | 0.63 | 0.65 | 0.72 |

It can be observed from Table 2 that for Paclitaxel, MOLI and DeepInsight-3D produced promising AUCs. For Cisplatin, Super.FELT had the highest, and for Gemcitabine (TCGA), MOLI produced the highest. For the remaining 5 drugs, Gemcitabine (PDX), Cetuximab, Erlotinib and Docetaxel, DeepInsight-3D produced the highest AUCs. The average AUC over all the seven datasets for the MOLI method was 0.63 and for Super.FELT was 0.65. DeepInsight-3D produced an encouraging average AUC of 0.72. For confusion matrix over seven datasets, please see Table S3 (Supplement File 1).

DeepInsight-3D can also perform feature selection via class-activation maps (CAMs) to identify genes of interest for each dataset. Since the data dimensionality is very large compared to the number of samples available, there is a high chance of producing an unstable model estimate. Furthermore, not all genes can be well represented in a limited pixel-framework. Appropriate feature selection would reveal background scientific mechanisms. Therefore, we applied an iterative way of conducting feature selection. Gene selection can be performed in 3 ways, 1) considering CAM values for every training sample, 2) taking an average of CAM over training samples, and 3) class-based CAM (described in the Methods section) where the average over a particular class is considered. In this work, class-based CAM has been applied for gene selection. Table 3 depicts the number of genes selected for each drug dataset. For parameters related to feature selection, see Table S4 and feature selection procedure in Figure S1. The activation maps are shown in Figure S2 and Figure S3.

**Table 3:** Gene selection using DeepInsight-3D

| Drug | #Genes | #Genes per class | CAM Threshold |
| --- | --- | --- | --- |
| Paclitaxel (PDX) | 1057 | [891, 520] | 0.25 |
| Gemcitabine (PDX) | 1108 | [882, 738] | 0.30 |
| Cetuximab (PDX) | 1229 | [1147, 692] | 0.25 |
| Erlotinib (PDX) | 1204 | [881, 1015] | 0.23 |
| Docetaxel (TCGA) | 1043 | [720, 576] | 0.25 |
| Cisplatin (TCGA) | 949 | [752, 278] | 0.25-0.30 |
| Gemcitabine (TCGA) | 1424 | [1034, 782] | 0.3 |

### Pathway-centric context of discovered gene sets

Gene sets identified as important by each drug-specific model were mapped to KEGG pathways and IPA knowledgebase, as described in the methods section. This analysis has revealed that the there was both a unique as well as a shared component that was identified as important for all of the drugs. In several cases most significantly enriched subsets have previously been reported in literature as being linked to particular drugs or that class of drugs. This suggests that the proposed system does have some functionality to not only improve quality of drug response prediction, but also allow discovery of meaningful biological processes that may be involved. Full results of these analyses are available as a Supplementary file XX; and relevant key findings are summarized below.

## Discussion

DeepInsight-3D extends the versatility of applying CNN to multi-layered tabular data. In this work, DeepInsight-3D provided very encouraging results on drug response multi-omics data. DeepInsight-3D was able to produce an average AUC of 0.72 over seven drug response datasets which is encouraging compared to competing methods in the literature.

Deep learning nets, such as CNN, have many merits, such as automatic feature extraction, finding hidden structures from hyper-dimensional data, finding higher-order statistics of image and non-linear correlations, economical use of neurons for large input sizes allowing much deeper networks are plausible with fewer parameters [29], and a parsimonious memory footprint. These properties of CNN can be integrated with the inception of DeepInsight-3D for non-image tabular data with multi layers.

In machine learning techniques for tabular data, any two features are considered mutually independent. However, DeepInsight tries to establish a relationship through the element arrangement step by positioning similar elements together and dissimilar ones apart [8]. DeepInsight-3D further extends this property to multi-layered data. Moreover, the application of DeepFeature is extended. DeepFeature enables a powerful means for the identification of biologically relevant gene sets and provides methodological basement for “explainable AI”. [9]. This has been integrated with DeepInsight-3D to simultaneously identify elements for multi layered data.

Although the results were promising, the severe scarcity of training and test samples hindered getting a reasonable model estimate. The same was true for MOLI and Super.FELT methods, as their results were sensitive to parameter tuning. In general, CNN works very well when the samples are sufficiently large. However, this was not the case in the work. Nonetheless, all these methods provided a good platform in this direction. DeepInsight-3D can perform sufficiently well when the sample size is sufficient such as in the case of single-cell analysis. This would be our future direction of work.

## Methods

This section covers the proposed DeepInsight-3D methodology. The model consists of the following constituents 1) image transformation by DeepInsight-3D, 2) ResNet-50 model of CNN architecture, 3) class-based CAM to find activation maps, and 4) element decoder to decode genes. These procedures are described hereunder.

### DeepInsight-3D: Conversion of multi-layered tabular data to image for CNN

Let a multi-layered sample be depicted by 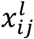, where *i* represents elements or features, *j* represents samples, and *l* represents layers. Therefore, a single layer data can be depicted as 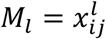 for *i* = 1,2, …, *d, j* = 1,2, …, *n* and *l* = 1,2, …, *L*, where *d* is the dimensionality of the data, *n* is the number of samples, and *L* is the total number of layers. For multi-omics data in this work, *L* = 3, which gives a multi-layered dataset *M* = {*M*_1_, *M*_2_, *M*_3_} ∈ ℝ^*d*×*n*×*L*^. The DeepInsight model [8] converts non-image data *M*_*l*_ to image data *E*_*l*_. The size of an image sample is *p* × *q*. The DeepInsight transform consists of dimensionality reduction techniques (such as t-SNE [30], UMAP [31] or Kernel PCA [32]), convex hull algorithm, rotation of Cartesian coordinates, finding pixel locations and mapping of elements to these pixel locations. We can obtain pixel locations by

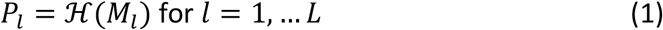

where *P*_*l*_ is the pixel locations of layer *l*, ℌ denotes the DeepInsight transform to find pixel locations and *M*_*l*_ is a single layer of training set (e.g. gene expression data). Once the framework of the locations is discovered using Eq (1), elements can be mapped to find the corresponding images, such as

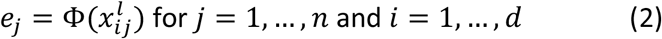

where Φ maps a non-image sample *x* ∈ ℝ^*d*^ to an image sample *e*_*j*_ ∈ ℱ^*p*×*q*^, here ℱ is a pixel-coordinates system, and, *p* and *q* are sizes of rows and columns, respectively. Therefore, from Eq (2) we get Φ: *x* → *e*. The transformation Φ also normalizes the values between [0,1] or [0,255]. In this work, norm-2 has been employed which was introduced in [8].

Thus, the first layer of image data (*l* = 1) obtained from Eq (2) is

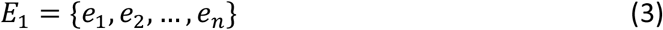

For simplicity, the superscript *l* is ignored on *e*_*j*_. However, this dataset obtained from Eq (3) is for layer *l* = 1. For *l* = 2, we did not compute the transform ℌ, however, only Eq (2) has been used to find *E*_2_. Similarly, for *l* = 3, we can obtain the dataset *E*_3_ from Eq (2). Therefore, for *l* = 1,‥, *L*, we get a multi-layered image dataset with common pixel locations *P*_1_. In this work, *L* = 3, so we get a 3D colored image of a multi-omics sample.

In the above model, it has been assumed that information from layer 1 is more than the other two layers, and that’s why all the other samples of the remaining two layers also mapped on *P*_1_. If it cannot be determined which layer has more information compared to others, then all the layers can be used simultaneously to find common pixel locations. In that case, transform ℌ for *l* = 1, …, *L* will be applied. However, it would produce multiple pixel locations (*P*_1_ … *P*_*L*_) and we need to find the common pixel locations from these pixel locations. This requires a two-stage process and is implemented in the DeepInsight-3D package by setting up the parameter *Parm*.*FeatureMap* to ‘0’. However, since this option has not been used in this work, the detailed description is avoided in this paper.

### CNN architecture for classification and feature selection

In this work, ResNet-50 has been used for CNN. For feature selection, we have incorporated class-activation maps (CAMs) [28]. However, other series nets supported by CAM can be used. ResNet-50 has a fixed input image size of 224 × 224 × 3. However, different image sizes can be used, as package resizes and corrects the size according to the requirements of the net. The last ReLu layer has been used to find activation maps. The activation maps express the region of interest for decision making. It provides 3 colored layers in order of importance as red, yellow and blue. Since the red zone is the most informative, it has been used for feature selection purposes using the element decoder (Fig. 1). The training set and validation set are used to estimate and validate the model. The test set is used to evaluate the performance of the trained model. For CAM, only the training set has to be used to compute activations. The default values of hypermeters of CNN net, such as momentum, L2 regularization and initial learning rate have been used (as per version 2 of the DeepInsight package https://alok-ai-lab.github.io/DeepInsight/). However, a Bayesian optimization technique has been employed for Cisplatin to tune the hyperparameters. Further description is given in the Supplement File.

### Class activation maps (CAMs) and element decoder

CAMs are computed for each image sample from the training set *e*_*j*_. CAM produce 3 colors and if we denote *R*_*j*_ as the computed CAM values of the red zone for a sample *e*_*j*_, then *R*_*j*_ > *threshold* depicts a region of interest for this sample. Since samples *e*_*j*_ falls in different classes (here respondents and non-respondents), we can take an average of *R*_*j*_ over the samples of a class. Therefore, class-based CAM can be computed as

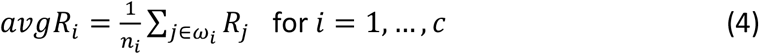

where *ω*_*i*_ denotes *i*-th class, *c* is the number of classes (here 2), and *n*_*i*_ is the number of training samples in this class.

For class-based CAMs, instead of taking *R*_*j*_ > *threshold*, one can consider *avgR*_*i*_ > *threshold* from Eq (4). Under this activated region, element decoder finds the gene subset. The decoder will locate the argument or index of a pixel falling under this region. A pixel *p*_*k*_, located at (*a*_*k*_, *b*_*k*_) is defined by normalized value [0,1]. However, depending upon the compression, it may contain one gene, more than one gene, or no gene. Searching all the pixels under the activated region (as defined by Eq (4)), would reveal a list of selected genes. This procedure will provide class-based features (or genes or elements), however, some elements could be common across different classes.

Let *G*_*i*_ be the gene subset found from the *i*-th class, then the overall selected genes are denoted as

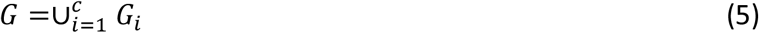

### Experimental setup

We used the same setup of datasets as done in [5], where training sets were collated from GDSC cell lines resource [2]. The test sets were collated from TCGA patients with the drug response [33] and PDX encyclopedia resource [6].

The data was downloaded from the Zenodo repository (https://zenodo.org/record/4036592) and correlated into the seven testing and training using R.

Test samples have two labels, non-responders (NR) and responders (R). The test set labels are exactly the same as [5] and are shown in Table 1. The total samples used for training models are also derived from GDSC resource, same as [5], however, the number of NR and R may be different. For all the training sets, first we applied a median of log *IC*50 to separate NR and R labels. This attempt balanced the NR and R samples in the training sets. However, in the case of Cisplatin, the validation accuracy was not promising, and so we then applied ‘mean’ to separate the labels.

Since the number of samples is very limited for all the drug response data, we augmented the training and validation sets during the training phase of CNN (see Methods for the details).

All the experiments were done on Intel Xeon Gold 5220R Server (2.2GHz) with 24 CPU cores and 2 parallel NVIDIA A100 PCIe GPUs (CUDA cores: 6912 with 40GB GPU memory on each A100 GPU). The operating system used was Linux (Ubuntu Desktop version 20.04).

### Pre-processing of mutation data

Cancer mutation data is most often extremely sparse, meaning that only a small number of different genes have consequential mutations in each sample. This presents a unique challenge when using it with a CNN classifier - as most inputs in this channel would be zero most of the time, it can result in inefficient use of information in that layer due to “dead” artificial neurons [34]. To counter this, we have used guilt-by-association principle to propagate the likely impact of mutations by using protein-protein interaction network. Briefly, the goal of this approach was to assign some part of an “impact” for each actual mutation to proximal genes in the network, as these are likely to be involved in similar biological functions. In this way, some meaningful value is assigned to each gene in all situations, while the information about actual mutations is still preserved by assigning them the highest possible score. Note that the fine calibration of the impact score is not necessary for this use-case, as neural network is able to discover its own optimal weighting as long as the generated distribution is consistent across all of the training set.

This was done by mapping all of the genes in the dataset to the corresponding proteins of the protein interaction network obtained from STRING database v11.0 [35]. A diffusion state distance matrix was calculated for the network based on the original definition of this distance metric [36]. Then, each node was assigned a score equal to the normalized inverse distance value of the closest mutated gene. In this way, the approach has facilitated the identification of possible functionally equivalent mutations as well as mutation hotspots, which have also been demonstrated to be an important network-based feature potentially predictive of clinical outcomes [37].

### Model evaluation

In order to validate DeepInsight-3D, the training sets (besides the test sets) were subdivided into two sets with a 90:10 ratio. The larger set was employed to estimate the model and the smaller set was applied to validate it. The AUC was computed on the test set. In general, the default parameters of DeepInsight (version 2) were employed (https://alok-ai-lab.github.io/DeepInsight/) for this method with a few variations (see Table S1, Supplement File 1 for details). Some important parameters were norm-2 normalization (log transform) [8], t-SNE to obtain a 2D plane for gene expression data, and that CNA and mutations were mapped to the 2D plane obtained by gene expression, as it is generally considered that gene expression has more information compared to the other profiles. For CNN, we applied a pre-trained ResNet-50. This transfer learning helped to achieve promising results. In order to have faster training, default parameters were applied for all the datasets, and the obtained performance was satisfactory. However, for Cisplatin, we did not get promising results. Therefore, for Cisplatin, Bayesian optimization technique of hyperparameter tuning was applied for ResNet-50. The hyperparameters that best performed over the validation set have been used for the test set (see Table S2, Supplement File 1 for details).

### Finding gene subsets through an iterative process

The number of genes in genomic or multi-omics data is typically very large, making it difficult to put all of them into a finite image size due to fixed technology limits. In this instance, quantized images are unavoidable, meaning that specific image pixels will carry several genes in a single spot. This leads to another issue of selecting a gene from those batch genes (where batch gene refers to a set of two or more genes having the same pixel location in the frame). To address this overlapping issue up to some extent, DeepInsight-3D can be run iteratively to gradually select the elements. The initial iteration will identify a subset of elements that can be utilized as input in subsequent iterations to find a smaller subset of genes or elements.

### Functional annotation and interpretation of identified gene sets

The analysis described above resulted in two gene lists (one each for responder and non-responder class) from each trained model that contained the genes identified as important for classifying training samples into a respective category. Functional interpretation of the recovered gene subsets was done individually, by mapping them onto metabolic and signaling pathways as defined by KEGG database [38]. This was followed by gene set enrichment analysis done using Fisher’s exact test with a Benjamini-Hochberg false discovery rate correction. A complementary perspective was produced using Ingenuity Pathway Analysis software from QIAGEN Digital Insights, which facilitates discovery of upstream/downstream regulatory context of particular genes and interpretation of likely effects on related mechanisms and biological processes.

## Conclusions

The proposed method, DeepInsight-3D demonstrates how data-to-image approach for analysis of biological data can effectively incorporate different types of ‘omics data and preserve the explicit connections between these layers by placing them in the same positions but in different channels of an input image. As was demonstrated in our previous work, once converted to image form data becomes suitable for use with image-specific convolutional neural network architectures. This study is the first to use this type of ‘omics integration and likewise the first to apply this type of approach to the problem of personalized cancer drug response prediction. Our results have shown that DeepInsight-3D can outperform previously proposed methods and can also be very powerful way to discover underlying important genes, which can then be interpreted understand the decisions made by the classifier and also identify key biological processes of potential interest.

## Acknowledgement

This work was funded by MEXT/JSPS KAKENHI Grant Number JP20H03240

## Author contributions

AS perceived, designed the classification and feature selection models, and wrote the first draft and contributed in the subsequent versions of the manuscript. AL designed the enrichment analysis, improving selection models and contributed in the first draft and thereafter of the manuscript. KAB build further the enrichment analysis, prepared the data, feature selection tools and helped in the manuscript writeup. TT perceived and contributed in the manuscript writeups. All authors read and approved the manuscript.

## Code Availability

DeepInsight3D software package (in Matlab), a dataset, installation instructions and user-manual are available from the GitHub link https://github.com/alok-ai-lab/DeepInsight3D_pkg. The example PDX_Paclitaxel dataset is also separately available from the link http://emu.src.riken.jp/DeepInsight/download_files/dataset1.mat, note the size is 88MB. The following links for other related packages can be accessed via http://www.alok-ai-lab.com/tools.php and/or http://emu.src.riken.jp/.

## Notes

https://github.com/alok-ai-lab/DeepInsight3D_pkg

http://emu.src.riken.jp/DeepInsight/download_files/dataset1.mat

http://www.alok-ai-lab.com/tools.php

http://emu.src.riken.jp/

### Competing Interest Statement

The authors have declared no competing interest.

### Summary of Updates

The authors' affiliations were not correct. Therefore, I revised it again. Apologies for the mistake. Kind regards Alok

