## Supplement-1 for "DeepInsight-3D for precision oncology: an improved anti-cancer drug response prediction from high-dimensional multi-omics data with convolutional neural networks"

### Supplementary File 1
### Feature selection and tuning of DeepInsight-3D parameters

#### 1. DeepInsight-3D parameters

The pipeline of DeepInsight-3D is based on the DeepInsight [1] architecture. The tabular data in a multi-layered form or multi-omics data are transformed to image samples for the application of CNN. It uses element arrangement using dimensionality reduction technique (DRT), followed by convex hull algorithm and rotation in the Cartesian coordinates. Thereafter, translated to pixel coordinates and mapping of original elements to these pixel locations. These steps are considered for a 3D matrix, and finally, a colored image is obtained for a non-image sample. The DRT options available for DeepInsight-3D are t-SNE [2], UMAP [3], kernel PCA [4] and PCA [5]. All possible distances of t-SNE can be employed, such as 'Euclidean', 'cosine', 'hamming', 'correlation', etc. In this work, 't-SNE' with 'Euclidean' distance has been used to find element locations for all the drug datasets. For t-SNE, if the number of elements or data dimensionality is above 5000, the Burnes-Hut algorithm is applied for faster processing; otherwise, the exact algorithm is used.

#### 2. CNN parameter

The ResNet-50 has been used for CNN implementation. The ResNet-50 configuration has a depth of 50. The hyperparameters such as learning rate, momentum, and L2regularization are identical for most drug datasets (except Cisplatin) and are given in Table S1. For the Cisplatin dataset, hyperparameters are selected using the Bayes optimization technique from a range of values. The description of these parameters is given in Table S2.

The hyperparameters (Table S1) are obtained by optimizing the validation error on one of the datasets, and the same values are used for all other datasets. These hyperparameters are set as default values in DeepInsight version 2 package:
https://github.com/alok-ai-lab/DeepInsight/tree/master/ver2

Table S1: ResNet-50 training options.

| *Variables* | *Values/range* |
|---|---|
| Net | ResNet-50 |
| Training option | Sgdm (Stochastic gradient descent with momentum) |
| InitialLearningRate | 4.98661e-5 |
| Momentum | 0.801033 |
| L2regularization | 1.25157e-2 |
| Max Objectives | 1 |
| MaxTime | 50x60x60 |
| Execution environment | multiple-GPU |
| MaxEpochs | 400 |
| Min batch size | 512 |
| Shuffle | Every-epoch |
| Augmentation | Yes |
| Image Size | 224 x 224 |

Table S2: ResNet-50 parameters options for Cisplatin dataset using Bayesian optimization technique. All the other parameters are same as Table S1.

| *Variables* | *Values/range* |
|---|---|
| InitialLearningRate | [1e-5 1e-1] |
| Momentum | [0.8 0.95] |
| L2regularization | [1e-10 1e-2] |
| Max Objectives | 10 |

The range of values was applied during the training session, and the best values were selected, which gave the least validation error.

Two types of norms are introduced in DeepInsight [1]. However, in this work, we applied Norm-2.

#### 3. Element versus feature

In this work, the term 'element' refers to the raw data, and 'feature' refers to the processed or extracted data. However, the term 'feature' encompasses 'element' since element becomes feature if identity transform is applied, i.e., $feature = identity \times element$.

The term 'element arrangement' in this paper is used because we utilized RNA-seq, CNA and somatic mutation data. However, this term can be considered identical to 'feature arrangement' such as protein sequences are converted to features vectors for certain applications.

#### 4. Feature mapping

The most informative layer out of the 3 layers is considered to derive pixel locations. Once the pixel locations are obtained, then three layers will be mapped onto these locations separately. This would give three 2D matrices. These matrices are used as R, G and B layers of a colored image sample. This way, a tabular three-layered sample (or multi-omics sample) is transformed to a colored image. The mapping of elements gives values or shades to locations. The element locations are determined by using the training set. If more than one element occupies the same spot, their averaged values are mapped onto this spot. For example, if locations of $g_i, g_j$ and $g_k$ are the same $(r, c)$ then $(g_i + g_j + g_k)/3$ will be mapped on this location. In this case, quantized compression will occur [6]. The validation and test sets used the same locations obtained from the training set.

The conversion of tabular sample to image gives an optimum pixel frame of size $p \times q$. The compression would be non-quantized if the desired frame size $m \times n$ is greater than $p \times q$ ($p \leq m$ and $q \leq n$). The compression can be quantized if $p > m$ and $q > n$. The quantized compression can be attained by adjusting the horizontal and vertical sizes of the pixel frame. In this case, it is possible that more than 2 elements can attain one location giving the average value of elements at this particular location. Therefore, as discussed in the previous example, if $k$ elements $\{g_1, g_2, \ldots, g_k\}$ having a common location, then average value $\frac{1}{k}\sum_{i=1}^{k} g_i$ will be mapped at this location. This is termed as the 'overlapping issue' in this work. It is desirable to minimize overlapping issue especially for element or feature selection process.

#### 5. Confusion matrix

The DeepInsight-3D produced an average AUC of 0.72 over seven drug datasets using test sets. The confusion matrices of test sets are depicted in Table S3.

Table S3: Confusion matrix on test sets produced by the DeepInsight-3D method.

| Method | PDX | PDX | PDX | PDX | TCGA | TCGA | TCGA |
|---|---|---|---|---|---|---|---|
| Drug | Paclitaxel | Gemcitabine | Cetuximab | Erlotinib | Docetaxel | Cisplatin | Gemcitabine |
| Confusion matrix | [24 14]<br>[2 3] | [10 8]<br>[2 5] | [28 27]<br>[0 5] | [13 5]<br>[1 2] | [6 2]<br>[4 4] | [1 5]<br>[14 46] | [10 26]<br>[3 18] |

#### 6. Feature selection using DeepInsight-3D

The activation of the multi-omics layers (gene expression, CNA and somatic mutation) is considered for finding genes. Since the number of genes is quite large, it was difficult to extract a meaningful subset in the first attempt due to the overlapping of elements on pixels. This problem was mitigated by applying the feature selection process in several iterations, as shown in Figure S1. In this case, activation maps produced by CNN are processed to the element decoder with a specified CAM threshold to obtain gene subsets (say $G_1$). The reorganized dataset corresponds with respect to gene subset $G_1$ will be processed back to the DeepInsight-3D model, which would give a smaller gene subset $G_2$. This process will continue until the desired number of genes is obtained. Therefore, this process will gradually reduce the features or elements so that genes selected due to overlapping of pixels can be reanalyzed. The term Stage-$j$ is used here to refer to the $j$-th iteration to obtain gene subset $G_j$. The iterative process is terminated when around 1000 genes are obtained.

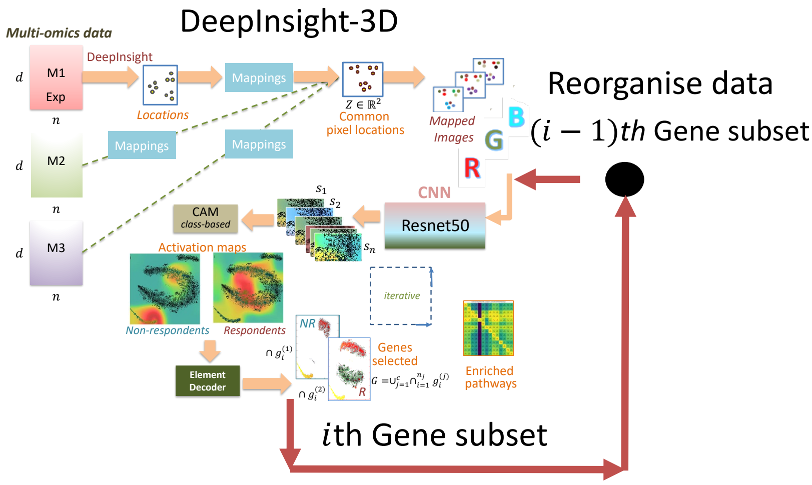


Figure S1: An iterative way of performing element or gene selection.

The details about gene subsets obtained, stages and CAM threshold are given in Table S4.

Table S4: Gene selection using DeepInsight-3D

| **Method**<br>**Drug** | **PDX**<br>**Paclitaxel** | **PDX**<br>**Gemcitabine** | **PDX**<br>**Cetuximab** | **PDX**<br>**Erlotinib** | **TCGA**<br>**Docetaxel** | **TCGA**<br>**Cisplatin** | **TCGA**<br>**Gemcitabine** |
|---|---|---|---|---|---|---|---|
| #Genes | 1057 | 1108 | 1229 | 1204 | 1043 | 949 | 1424 |
| Stage | 5 | 4 | 6 | 4 | 6 | 3 | 5 |
| CAM Threshold | 0.25 | 0.3 | 0.25 | 0.23 | 0.25 | 0.3@Stages1-2<br>0.25@Stage3 | 0.3 |

From the table, for the PDX Paclitaxel dataset (2nd column), DeepInsight-3D was run up to Stage-5 (i.e., five iterations), and the CAM threshold for each of the stages was 0.25. For TCGA Cisplatin, the CAM threshold for Stage 1 and Stage 2 was 0.3, and for Stage 3 was 0.25. The number of total genes selected per dataset is also highlighted in green color.

Figure S2 shows the activations of the PDX Paclitaxel dataset at Stage 1 and Stage 5 (last stage). Only a training set of Paclitaxel has been used to find the activations. The selected genes are also depicted on the right-hand side of Figure S2.

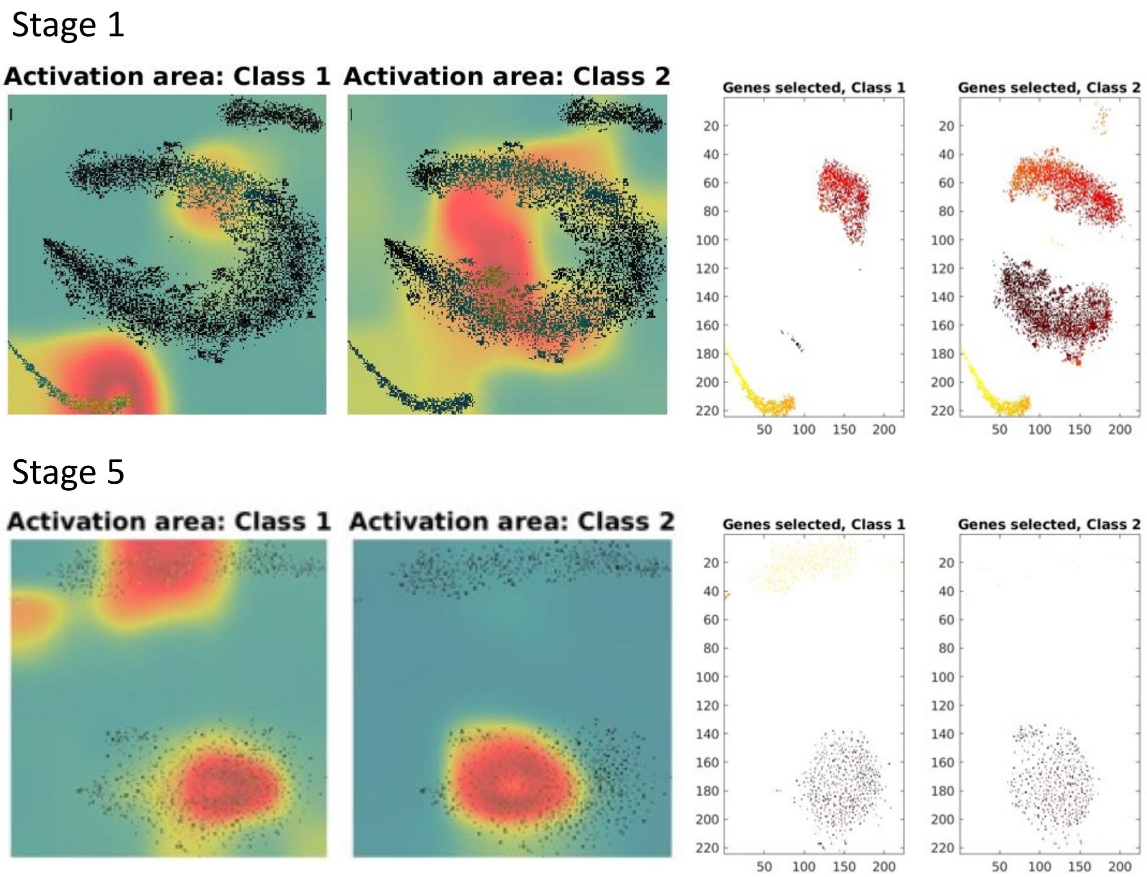


Figure S2: Activations and gene selection on Paclitaxel at Stage 1 and Stage 5. Class 1 is non-respondents and Class 2 is respondents. The left side of the figure depicts the activations in Class 1 and Class 2, and the right side illustrates the selected genes in the respective classes.

Similarly, for all the drug datasets, the activations are shown in Figure S3 for the last stages (see Table S4 for details about stages in different datasets). The preferential order of activation is from red to yellow to blue; i.e., the genes under the red activation zone are the most desirable. We have only considered genes corresponding to the red activation and discarded the other regions such as yellow, blue and without activation. As mentioned in the manuscript, class-based CAM has been adopted for element selection. Therefore, only two activation maps are obtained (since only two classes are present in the dataset) and from each map, a subset of genes can be derived specific to ‘responders’ and ‘non-responders’. It can be observed by comparing the activation maps of Class 1 and Class 2 from Figure S3 that many genes are common in both the classes. However, some genes are different, which helps in identifying different behavior toward a given cancer drug. Therefore, the DeepInsight-3D method finds genes which are representative and, at the same time, different responses.

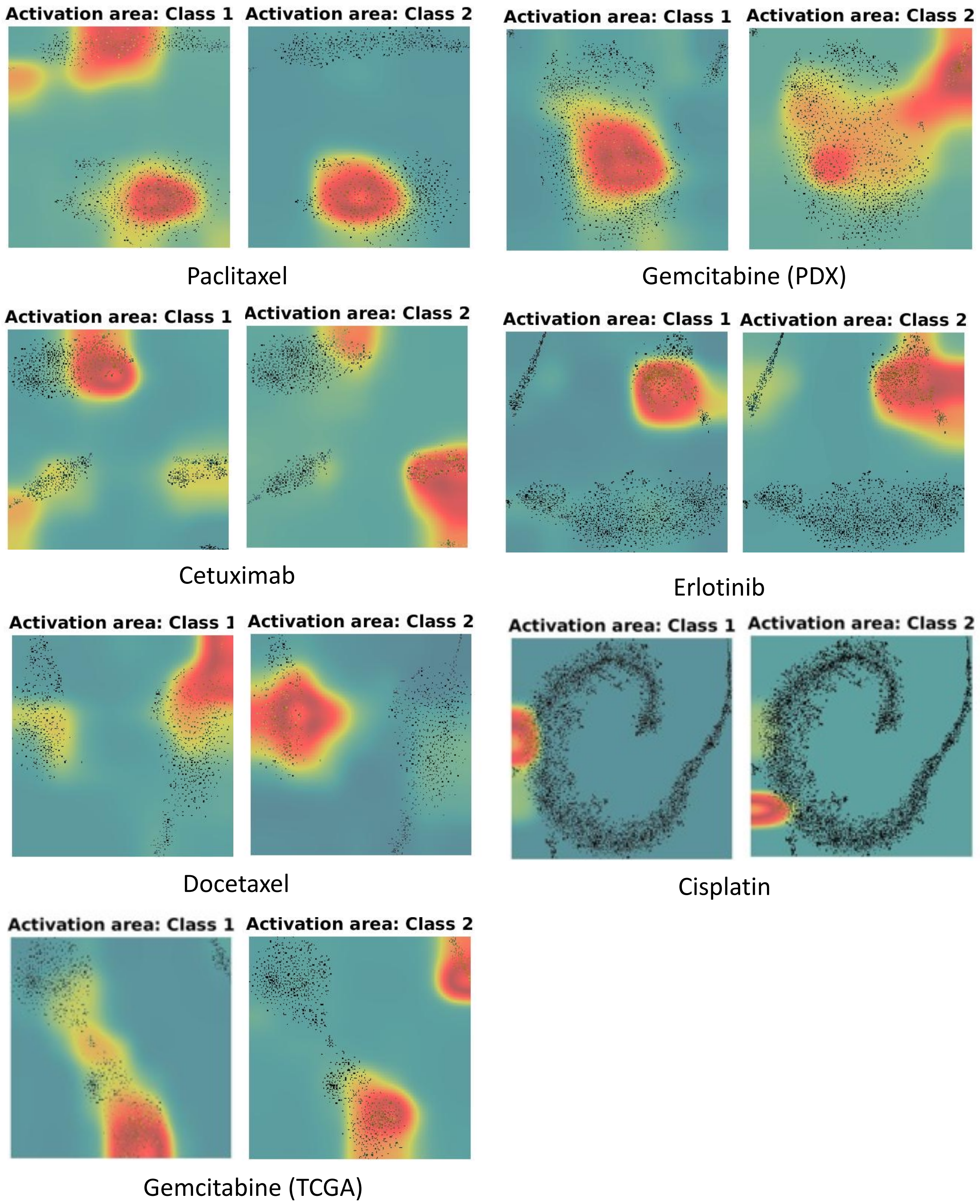


Figure S3: Class activations for seven datasets used in this study where Class 1 is 'non-responders' and Class 2 is 'responders'.
